# High genomic stability of *w*Mel *Wolbachia* after introgression in three geographically distinct *Aedes aegypti* populations

**DOI:** 10.1101/2025.02.24.638572

**Authors:** Kimberley R. Dainty, Jane Hawkey, Endah Supriyati, Louise M. Judd, Alexander Uribe, Iván D. Vélez, Dang D. Anh, Eggi Arguni, Warsito Tantowijoyo, Scott L. O’Neill, Kathryn E. Holt, Cameron P. Simmons, Heather A. Flores

## Abstract

The introgression of *w*Mel *Wolbachia* into *Aedes aegypti* populations is being used for the biocontrol of arboviruses such as dengue and chikungunya in 15 countries to date. A *w*Mel infection in *Ae. aegypti* both reduces the transmission of viruses by the mosquito and causes a reproductive manipulation that aids *w*Mel introgression into naïve populations. However, a critical concern is whether *w*Mel could evolve over time, potentially diminishing these desired phenotypes. Here, we investigated the stability of the *w*Mel genome in *Ae. aegypti* released for biocontrol in Colombia, Indonesia, and Vietnam. We sequenced the *w*Mel genome at the start of releases and up to six years after *w*Mel introgression into each population. Our study identifies very few genomic changes, suggesting the *w*Mel genome is not rapidly evolving despite its release into three geographically different field sites and subsequent exposure to novel environments. These results align with previous *w*Mel sequencing studies from Australia and provide strong evidence for the long-term genomic stability of *w*Mel, reinforcing its potential as a reliable biocontrol tool against *Ae. aegypti*-transmitted arboviruses.

## Introduction

Human pathogenic viruses such as dengue and Zika are spread primarily by *Aedes aegypti*. The burden of these diseases has increased substantially over the past 50 years due to increasing globalisation, the rise in population-dense cities, and expansion of the geographical breeding range of *Ae. aegypti* (Gubler 2002; Bonizzoni et al. 2013; Mayer et al. 2017). To reduce the burden of these viruses, a public health intervention utilising the bacterial endosymbiont, *Wolbachia pipientis* (*w*Mel strain), has been deployed in 15 countries to date (World Mosquito Program, n.d.).

Stable *w*Mel infection of *Ae. aegypti* reduces the ability of the mosquito to transmit multiple human pathogenic viruses such as dengue, chikungunya and Zika (Aliota, Peinado, et al., 2016; Aliota, Walker, et al., 2016; Carrington et al., 2018; Pinto et al., 2021; Rocha et al., 2019; Tan et al., 2017; Utarini et al., 2021; van den Hurk et al., 2012; Ivan Dario Velez et al., 2023; Walker et al., 2011). Effective biocontrol of these pathogens requires the spread and persistence of a *w*Mel infection in *Ae. aegypti*. This is facilitated by both *w*Mel maternal transmission, and the *w*Mel-induced reproductive modification, cytoplasmic incompatibility (CI), which aids the introgression of *w*Mel through naïve populations (Ant et al., 2018; Walker et al., 2011).

With an endorsement from the World Health Organisation and recent investment of funds geared towards expansion of this biocontrol method (World Health Organisation, 2025; Oxitec, 2025), it is likely that the next decade will see further global deployment of *w*Mel and other *Wolbachia* strains for use in *Ae. aegypti* and other mosquito species to reduce their disease burdens. One risk of using an endosymbiont as a biocontrol tool is the potential for this endosymbiont to undergo genomic changes which limit the phenotypic effects the endosymbiont has on its host. In this instance, the cessation of virus inhibition or the ability for *w*Mel to spread through or remain sustained in field populations would halt the effectiveness of the biocontrol. The genomic variation of *w*Mel has been examined in Australian *Ae. aegypti*. Studies compared the *w*Mel genome from *Ae. aegypti* in Cairns, Australia when first released and up to nine years post release to that of the reference *w*Mel genome sequenced directly from *D. melanogaster* (Wu et al., 2004). These studies found very little genomic variation after transinfection and field establishment, suggesting the *w*Mel genome is stable in Australian *Ae. aegypti* (Dainty et al., 2021; Huang et al., 2020; Ross et al., 2022).

After the success of Australian releases, *w*Mel was introgressed into *Ae. aegypti* populations in a further 14 countries, including Indonesia, Vietnam, and Colombia (Hien et al., 2021; Tantowijoyo et al., 2020; Iván Darío Velez et al., 2023). Exposure of *w*Mel to geographically diverse environments in their novel *Ae. aegypti* host again prompted concerns this would result in the local adaptation of *w*Mel. Studies of *w*Mel in *D. melanogaster* have identified strong geographical haplotypes (Early & Clark, 2013; Richardson et al., 2012) which have been hypothesised to be influenced by selection and local adaptation rather than genetic drift alone (Early & Clark, 2013). To date there is no conclusive field data indicating the crucial phenotypic effects *w*Mel induces in *Ae. aegypti* have been impacted by *w*Mel evolution. However, *w*Mel introgression and stability in fields sites has been highly variable. While many operational factors related to field releases such as number, life stage and method of mosquitoes released as well as local climates and host fitness may underlie this variation, changes to the *w*Mel genome may also explain the variability observed. One way to investigate this and the future potential for this to occur is to understand the breadth and spread of genomic changes occurring to the *w*Mel genome when used as a biocontrol agent. Here we examined the genomic changes to the *w*Mel genome after introgression into three geographically and environmentally diverse *Ae. aegypti* populations in Indonesia (first released January 2014), Vietnam (first released May 2014), and Colombia (first released June 2015). Overall, we found that the *w*Mel genome is highly stable in the 4-6 years of sampling post release in each population, suggesting the use of this endosymbiont as a biocontrol agent is not inducing rapid evolution that may lead to the loss of critical phenotype effects.

## Methods

### Sample collection and processing

#### Colombian mosquitoes

Colombian mosquito samples used for sequencing were collected from releases performed in the París comuna of Bello, Antioquia. Pilot releases in the central part of París took place between June and December of 2015, and a phase 2 release throughout the remaining París comuna took place between July and August 2016 (Iván Darío Velez et al., 2023). Female *w*Mel-infected *Ae. aegypti* mosquitoes were collected on March 19^th^ 2015 from an outbred pre-release colony which was used for subsequent field releases that commenced in June 2015. Ten individual mosquitoes were chosen for sequencing, and *w*Mel genomes represent the 2015 early-release *w*Mel genomes from Colombia. Female *w*Mel-infected *Ae. aegypti* mosquitoes were collected from the pilot release region of París on October 29^th^ 2019 via BG Sentinel traps (Biogents AG, Germany). These 24 *w*Mel genomes sequenced were from mosquitoes collected across 5 individual BG Sentinel traps and represent the 2019 recent-collection *w*Mel genomes from Colombia.

#### Indonesian mosquitoes

Indonesian mosquito samples used for sequencing were collected from the Sleman district of Yogyakarta, where *w*Mel releases took place between January-June 2014 (Tantowijoyo et al., 2020). Female *w*Mel-infected *Ae. aegypti* mosquitoes were collected from Sleman district on February 23^rd^ 2015 via BG Sentinel traps. These 9 *w*Mel genomes sequenced were from mosquitoes collected from 9 individual BG Sentinel traps and represent the 2015 early-release *w*Mel genomes from Indonesia. Female *w*Mel-infected *Ae. aegypti* mosquitoes were collected again from this same district on February 20^th^ 2020 via BG Sentinel traps. These 30 *w*Mel genomes sequenced were from mosquitoes collected across 26 individual BG Sentinel traps and represent the 2020 recent-collection *w*Mel genomes from Indonesia.

#### Vietnamese mosquitoes

Vietnamese mosquito samples used for sequencing were collected from Tri Nguyen Island, where *w*Mel releases took place over a 27-week period from May to November 2014 (Hien et al., 2021). Female *w*Mel-infected *Ae. aegypti* mosquitoes were collected from the central and southern regions (hamlets 2 and 3) of Tri Nguyen Island (see Figure 1, Hien et al., 2021; Figure 1A, Lee et al., 2022), via BG Sentinel traps between the 27^th^ October 2015 and the 3^rd^ September 2015. Mosquitoes were collected exclusively from the central and southern end of Tri Nguyen Island as this was where *w*Mel infection prevalence was highest. These 9 *w*Mel genomes sequenced were from mosquitoes collected from 8 individual BG Sentinel traps and represent the 2015 early-release *w*Mel genomes from Vietnam. Female *w*Mel-infected *Ae. aegypti* mosquitoes were collected from the southern end of Tri Nguyen Island where *w*Mel was still established via BG Sentinel traps between the 28^th^ May 2019 and the 4^th^ June 2019. These 18 *w*Mel genomes sequenced were from mosquitoes collected from 6 individual BG Sentinel traps and represent the 2019 recent-collection *w*Mel genomes from Vietnam.

**Figure 1.**
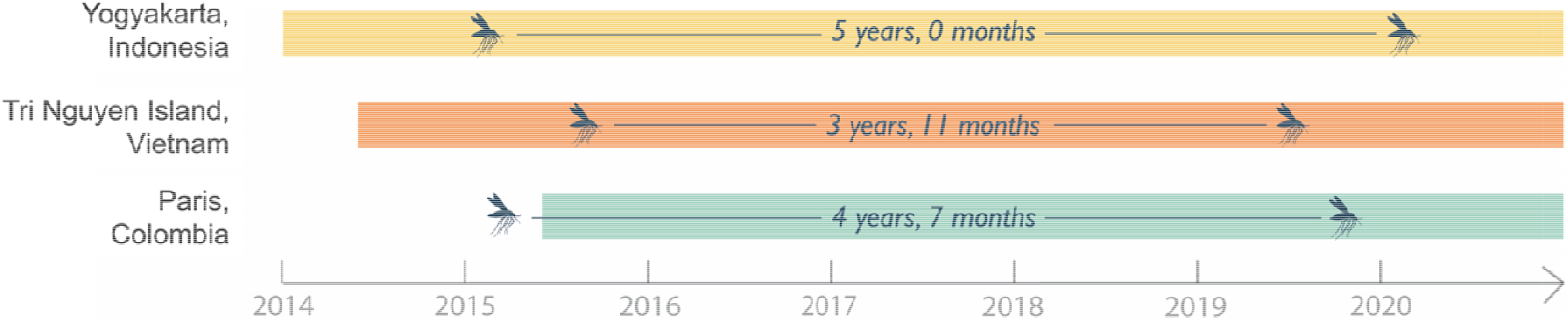
Timeline of the *w*Mel genome sampling. Release and establishment of *w*Mel-infected *Ae. aegypti* in Indonesia, Vietnam, and Colombia are indicated by yellow, orange, and green bars respectively, with the start of each colour block indicating initiation of *w*Mel-infected mosquito releases. Time points at which *w*Mel-infected *Ae. aegypti* were collected for sampling from each site are indicated by black mosquitoes. Time periods between collection times are displayed in black text.

#### All mosquitoes

DNA was extracted from the abdomens of single mosquitoes and used for sequencing as this would enrich for *w*Mel which is at higher density in the ovaries and Malpighian tubules (Amuzu & McGraw, 2016; Fraser et al., 2020). Sequencing libraries were performed from single DNA extracts, and thus, each genome represents the diversity present within an individual mosquito. Single abdomens were homogenised using a hand-pestle in Buffer ATL before DNA was extracted using the MagAttract HMW DNA Kit (QIAGEN) with a 50 μL elution in nuclease-free water following manufacturer’s instructions. Libraries were generated for each single mosquito sample using the Illumina DNA Prep (M) Tagmentation kit (Illumina®) using unique index tags. All Illumina DNA Prep libraries were made according to manufacturer’s directions, excepting all reactions were scaled to 25% of recommended volumes and 25ng of template DNA was used. Samples were sequenced via the Illumina platform using NovaSeq 6000 S4 Reagent Kit v1.5 (300 cycles) (Illumina®), generating 150-base-paired-end reads.

### Analysis

Illumina reads belonging to *Ae. aegypti* (accession NC_035107.1) were identified using BioBloom Tools v2.3.2 (Chu et al., 2014) using default parameters and excluded from downstream analysis. The remaining Illumina reads, not trimmed before analysis, were mapped to the *D. melanogaster w*Mel reference genome (Wu et al., 2004) (NC_002978.6) and single nucleotide variants identified using the RedDog V1b.11 pipeline (https://github.com/katholt/reddog) using standard parameters and following the developers’ guidelines. Briefly, this pipeline maps reads to the reference sequence using Bowtie2 version 2.2.3 (Langmead & Salzberg, 2012) before SNPs with QUAL values ε 30 are called with SAMtools version 1.9 (Li, 2011). Mean Illumina read depth for Colombian samples was 262.49x, for Indonesian samples was 846.51x, and for Vietnamese samples was 61.63x. Coverage was 100% for all samples. Gene copy number variation was analysed by normalising the sequencing depth of coverage of each gene in sequenced genomes by dividing the mean sequence depth for each gene by the mean sequence depth of the whole genome. Details regarding samples and associated sequencing data are available in Supplemental Data File 1.

Low frequency variants were detected using LoFreq V2.1.3.1 (Wilm et al., 2012) with default parameters. Variants were removed from the final data set if strand bias was ε 10. Variants were annotated with SnpEff V5.2, using the classical style of annotation and the database *Wolbachia* endosymbiont of *Drosophila melanogaster* GCA 000008025 (Cingolani et al., 2012). In cases where multiple annotations were identified for a SNP, the first annotation listed was used for analysis. Heatmaps were generated with ggplot2 (v3.5.1) (Wickham, 2016) using RStudio (v2024.12.0.467).

Insertion sequence elements were identified by using the *w*Mel reference genome (NC_002978.6) to search against the ISfinder (Siguier et al., 2006) database via the ISsaga (Varani et al., 2011) web server available at http://issaga.biotoul.fr/issaga_index.php. IS queries that were identified to have greater than 80% similarity (Supplementary Table 3) were mapped to the short-read sequencing data for each sample using ISMapper (Hawkey et al., 2015) to identify IS insertion sites with default parameters. Insertion or deletion of IS elements identified as imprecise (*) or uncertain (?) were considered mapping artefacts.

## Results

*No novel genomic changes occurred in the establishment of country-specific release lines* We sequenced *w*Mel from 9 Indonesian (Sleman district), 9 Vietnamese (Tri Nguyen Island), and 10 Colombian (París) individual mosquitoes collected just prior to or soon after introgression of *w*Mel (Figure 1). To generate these release strains, *w*Mel-infected Australian *Ae. aegypti* mothers were backcrossed to *Ae. aegypti* males from Indonesian, Vietnamese, and Colombian populations. *w*Mel from Australian *Ae. aegypti* used in these backcrosses has previously been shown to differ by only one single nucleotide polymorphism (SNP) different from the *D. melanogaster w*Mel reference sequence (NC_002978.6) (Dainty et al., 2021). Therefore, we compared their genomes to the *D. melanogaster w*Mel reference sequence (NC_002978.6) to identify changes that occurred during the establishment of country-specific release lines.

Across all pre/early release samples, only a single SNP was identified. We identified SNP_1174712 in 4/10 *w*Mel genomes collected from Colombia in 2015 (Figure 2). This SNP causes a synonymous change of base T->C in the WD_1228 gene, the product of which is a hypothetical protein (Table 1). Previously, this SNP has been identified in *w*Mel from Australian *Ae. aegypti* in up to 30% of individuals (Dainty et al., 2021; Huang et al., 2020; Ross et al., 2022), suggesting it was likely introduced into the Colombian background during backcrossing. SNP_1174712 was not found in the Indonesian or Vietnamese populations. These two *w*Mel-infected *Ae. aegypti* strains were generated via backcrossing of the same Australian population known to carry the SNP. Since this SNP was not fixed in the Australian *w*Mel population, it is possible this SNP was not inherited by the final Indonesian and Vietnamese colonies or inherited in low prevalence such that it was not sampled. Alternatively, we can’t discount the possibility that this SNP may cause a fitness disadvantage in these populations and thus has been selected against prior to the first collections and sequencing of these populations.

**Table 1.**
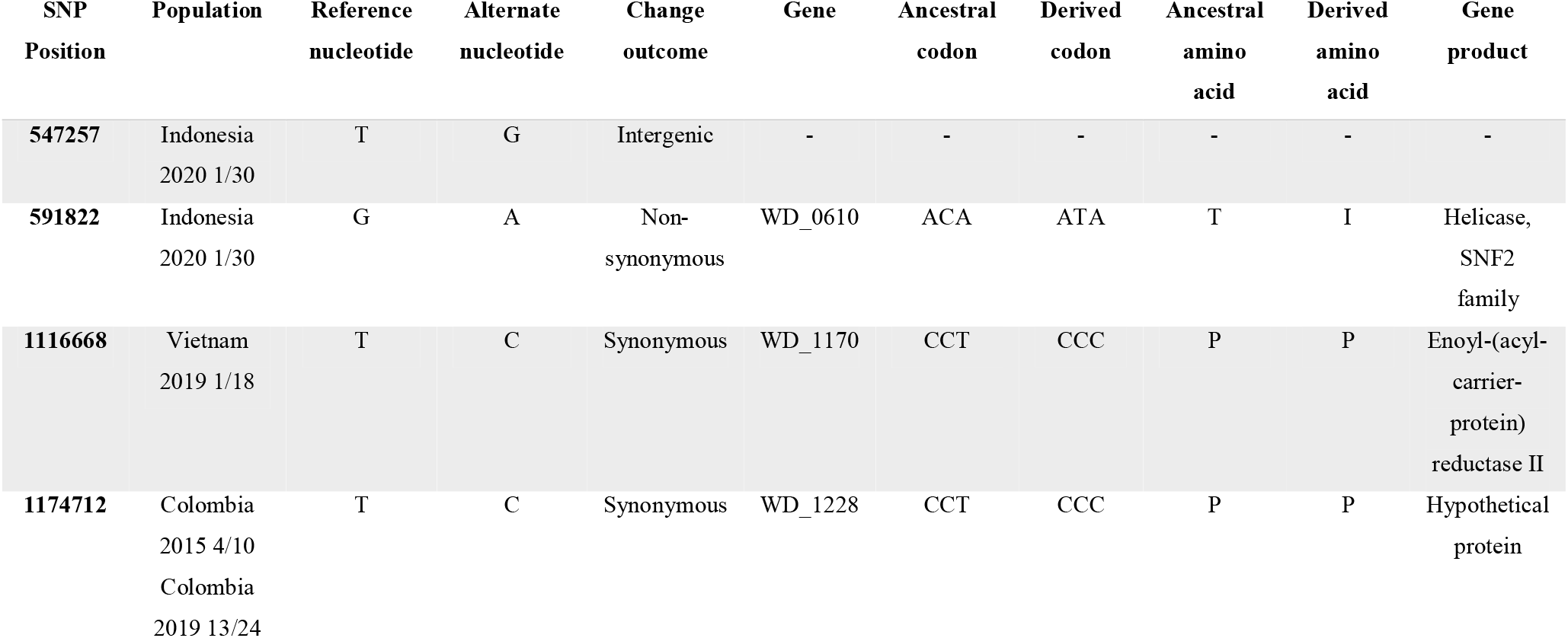
SNPs present in *w*Mel *Wolbachia* sequenced from *Ae. aegypti* collected in Colombia, Indonesia, and Vietnam compared to the *D. melanogaster w*Mel reference sequence.

**Figure 2.**
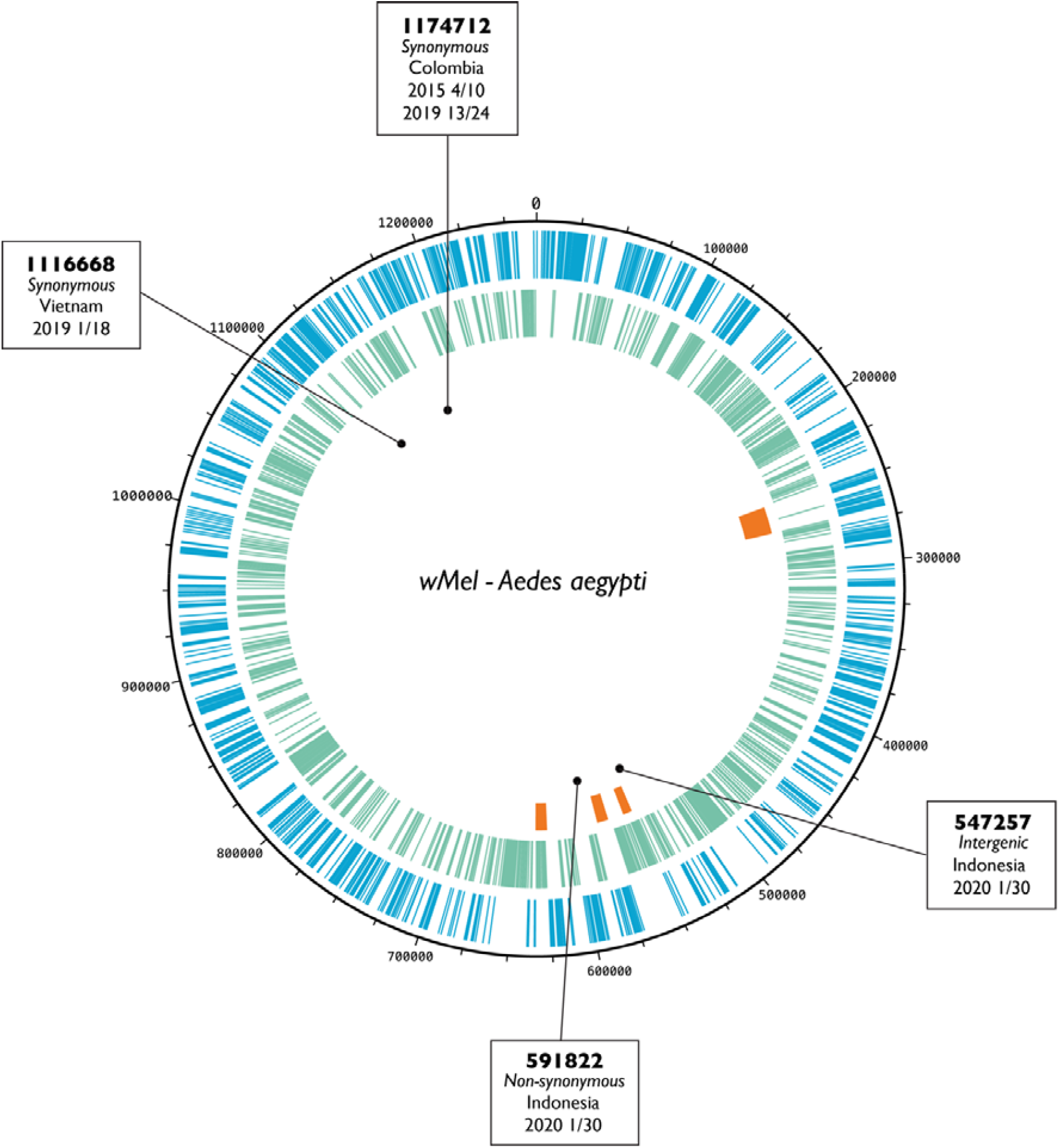
SNPs present in *w*Mel *Wolbachia* collected from *Ae. aegypti* compared to the *w*Mel reference sequence from *D. melanogaster*. Colours corresponding to the following: blue bars represent forward strand gene, green bars represent reverse strand genes, orange segments represent phage regions (in clockwise order WO-A, pyocin-like, WO-B, WO-B. SNPs are represented by black dots with corresponding information in connected boxes. Box information includes SNP position, mutation outcome, and population and number of genomes that contained the SNP.

### wMel genome is highly stable up to 6 years post-field release across all three field sites

We next sequenced *w*Mel genomes from Indonesian, Vietnamese, and Colombian *Ae. aegypti* populations 4-6 years post-release, to determine genomic changes influenced by field establishment and differing environments. Across the *w*Mel genomes sequenced here (30 from Indonesia, 18 from Vietnam, 24 from Colombia), one recurrent and three unique SNPs were identified in comparison to early field caught mosquitoes. Of these three unique SNPs, none were present in across multiple locations, and all three were found only once.

SNP_1174712 was again observed only in Colombian mosquitoes. Similar to Australian populations, we again observed that this SNP did not significantly increase nor decrease in frequency (4/10 to 13/24), suggesting it is not under selection in the Colombian population. SNP_547257 was found in one *w*Mel genome collected from an Indonesian *Ae. aegypti* in 2020 (Figure 2). This SNP causes an intergenic change of T->G at position 547257 (Table1) and is thus unlikely to be functionally impactful. Next, SNP_591822 present in one *w*Mel genome collected from an Indonesian *Ae. aegypti* in 2020 causes a non-synonymous change at position 591822 in the genome (Figure 2). This SNP constitutes a base change from G->A resulting in an amino acid change from threonine to isoleucine in gene WD_0610, the product of which is a SNF2 family helicase (Table1). This family of proteins have been shown to be responsible for directing energy from ATP hydrolysis into the remodelling of chromatin structures as well as processing of DNA damage repair, and transcriptional replication (Eisen et al., 1995; D. P. Ryan & Owen-Hughes, 2011). As this SNP was only found in 1/30 genomes sequenced from Indonesia in 2020, this indicates either this SNP has been of recent occurrence, or it is providing no evolutionary benefit and thus not spreading throughout the population. Lastly, SNP_1116668 was found in one *w*Mel genome from *Ae. aegypti* collected in Vietnam in 2019 (Figure 2). This SNP causes a synonymous base change from T->C in the gene WD_1170 which produces an enoyl-(acyl-carrier-protein) reductase II product (Table1). Again, as this SNP was found in a single genome only, and is causing a synonymous change, it is unlikely to impact the gene function.

In addition to these three polymorphisms and the previously described SNP_1174712, one further SNP was identified in all 100 genomes sequenced when compared to the *w*Mel reference genome, a SNP at position 1097797. However, this SNP has previously been shown to likely be an error in the *D. melanogaster w*Mel reference sequence (Dainty et al., 2021) rather than a novel genomic change, and is thus not included here. Additionally, when compared to the *D. melanogaster w*Mel reference genome, seven indels were identified in the genomes sequenced here. These seven indels were previously described in the *Ae. aegypti w*Mel genomes sequenced in (Dainty et al., 2021), and again shown to be errors in the *D. melanogaster w*Mel reference sequence.

Gene copy number variation was analysed in each genome by normalising gene mean sequencing depth to the mean depth coverage of the entire genome. No regions of the genome showed distinct copy number variation in any of the genomes (Supplementary Figure 1), albeit many early collection samples showed high variation in coverage levels possibly due to degradation of sample quality over time. The sequenced genomes were also analysed to assess insertion sequence (IS) movement, using the ISsaga and ISMapper programs. However, no evidence for movement, deletion, or novel insertion of any IS element was identified in any of the genomes. Due to the limitations of short-read sequencing, large scale structural changes cannot be ruled out in these genomes. However, given the lack of identified SNPs and indels, and the previously documented stability of the *w*Mel genome in Australian *Ae. aegypti*, it is unlikely that these genomes have undergone such structural changes.

### Evidence of low frequency wMel genomic variation within a mosquito

The above results looked at fixed SNPs of *w*Mel genomes at the population level across individual *Ae. aegypti* females. However, our sequencing was performed on single female samples allowing us to investigate *w*Mel variants *within* a single mosquito. We first investigated whether the four fixed SNPs identified in this study were present at low frequency across other samples. We found that SNPs 591822, 1116668, and 1174712 were only present in the single samples identified above. However, SNP 547257 was present at lower frequency in one additional Indonesian sample from 2015 (Supplemental Figure 2). We next compared low frequency variants across each population. We found a number of low frequency variants within each individual mosquito; however, the majority of these were at frequencies less than 10% (Supplementary Table 1). Averaged across each collected population, the number of variants with frequencies between 10-20% varies between 38.17 and 157.5 variants per genome. The average number of variants with frequencies above 20% ranges between 3.17 and 50.33 variants per genome (Table 2). The number of variants did not correlate with either *w*Mel density or sequencing depth (Supplementary Table 2). Interestingly, the average level of *w*Mel variation within a mosquito decreased in each population in later collections despite many generations in the field, including the prevalence of missense mutations which are more likely to have functional impacts for selection to act upon. Collectively, this suggests there are low levels of standing variation in these genomes that have not substantially increased in frequency nor yet fixed in the population.

**Table 2.** Mean number of low-frequency SNPs at frequencies of 10-20% and >20% identified per genome from each population. Mean number of variants predicted to cause missense, synonymous, or other impacts for each frequency is also reported.

| Population Collection (Sample size) | Mean # of SNPs at 10-20% | Mean # of SNPs with predicated impact variants at 10-20% (%) |  | Mean # of SNPs >20% | Mean # of SNPs with predicated impact variants at >20% (%) |  |
| --- | --- | --- | --- | --- | --- | --- |
| Indonesia 2015 (9) | 85.78 | Missense | 10.88 | 5.33 | Missense | 20.83 |
|  |  | Synonymous | 16.71 |  | Synonymous | 14.58 |
|  |  | Stop codon | 0.78 |  | Stop codon | 0.00 |
|  |  | Other | 71.63 |  | Other | 64.58 |
| Indonesia 2020 (30) | 68.73 | Missense | 12.37 | 5.03 | Missense | 9.93 |
|  |  | Synonymous | 14.26 |  | Synonymous | 4.64 |
|  |  | Stop codon | 0.68 |  | Stop codon | 0.00 |
|  |  | Other | 72.70 |  | Other | 85.43 |
| **Vietnam 2015 (9)** | 139.67 | Missense | 19.57 | 50.33 | Missense | 15.01 |
|  |  | Synonymous | 35.08 |  | Synonymous | 22.96 |
|  |  | Stop codon | 0.48 |  | Stop codon | 0.44 |
|  |  | Other | 44.87 |  | Other | 61.59 |
| **Vietnam 2019 (18)** | 44.5 | Missense | 12.36 | 15.94 | Missense | 10.10 |
|  |  | Synonymous | 23.10 |  | Synonymous | 8.01 |
|  |  | Stop codon | 0.12 |  | Stop codon | 0.35 |
|  |  | Other | 64.42 |  | Other | 81.53 |
| **Colombia 2015 (10)** | 157.5 | Missense | 15.81 | 26.1 | Missense | 8.12 |
|  |  | Synonymous | 22.03 |  | Synonymous | 8.43 |
|  |  | Stop codon | 0.70 |  | Stop codon | 0.00 |
|  |  | Other | 61.46 |  | Other | 82.76 |
| **Colombia 2019 (24)** | 38.17 | Missense | 13.97 | 3.17 | Missense | 6.58 |
|  |  | Synonymous | 20.63 |  | Synonymous | 22.37 |
|  |  | Stop codon | 0.55 |  | Stop codon | 0.00 |
|  |  | Other | 64.85 |  | Other | 71.05 |

## Discussion

*Wolbachia* biocontrol is transforming the fight against *Ae. aegypti-*transmitted arboviruses with the *w*Mel strain being deployed across diverse geographic and ecological landscapes. Given the evolutionary pressures imposed by these novel environments, we hypothesized that *w*Mel would be undergoing adaptive genomic changes, which could potentially explain variation in *w*Mel introgression and stability observed across field sites. Our study however identified only minor genomic changes present at low frequency in the three populations studied, thus suggesting that despite environmental pressures and novel hosts, the *w*Mel genome remains remarkably resilient, reinforcing its robustness as a long-term biocontrol strategy.

The evolutionary rate of *w*Mel in its native host *D. melanogaster* has been estimated to be low compared to other bacteria (Duchêne et al., 2016), at 6.8×10^-10^ substitutions per site per *D. melanogaster* generation (Early & Clark, 2013; Richardson et al., 2012). Assuming this evolutionary rate is similar in *Ae. aegypti*, combined with the relatively short time *w*Mel has been established in these populations (∼5 years, 12 generations per year), this rate predicts approximately 0.05 substitutions to have occurred per genome. Thus, the level of variation reported here is consistent with this estimation suggesting *w*Mel is evolving similar to levels observed in its natural host.

Although low levels of fixed SNPs were identified in the studied *w*Mel genomes, we did identify higher levels of low frequency polymorphisms within *w*Mel genomes from individual mosquitoes, indicating there is mutational input to these genomes. Previous *w*Mel sequencing from Australian mosquitoes was performed on pooled samples (Dainty et al., 2021). Our sequencing here was performed on single female mosquitoes allowing us to attribute this variation to heteroplasmy of *w*Mel within single mosquitoes. Our analysis shows that on average 86.6 variants occur per genome (with frequency >10%), with this number reaching more than 400 in some genomes. Clearly, however, the prevalence of these polymorphisms are not showing significant increases over time nor being transmitted at high levels, as evidenced by the low levels of SNP fixation detailed in this study. Severe bottlenecks such as those that occur during maternal transmission have been suggested to contribute to evolution of insect endosymbionts by altering the efficacy of purifying selection (Moran, 1996; Wernegreen, 2002). Additionally, our sequencing included *w*Mel found in somatic tissues within the abdomen, a dead end for this variation, as *Wolbachia* is maternally transmitted through the germline. Our data however suggest there are reasonable amounts of standing genetic variation present in *w*Mel populations that may allow *w*Mel to respond to various selective pressures it may experience in its mosquito host and/or environment.

These findings in combination with previous reports (Dainty et al., 2021; Huang et al., 2020; Ross et al., 2022), show the *w*Mel genome to be stable in four geographically distinct *Ae. aegypti* populations. Stability of the desired *w*Mel-induced phenotypes is critical for the success of the *Wolbachia* biocontrol methods, thus these findings in combination with entomological studies reflect positively on its continued success (Ary A. Hoffmann et al., 2014; Ross et al., 2022). However, *w*Mel is only one element of the tripartite interaction in this biocontrol method. In addition to *Wolbachia*, the *Ae. aegypti* host or the viruses that *Wolbachia* inhibit may evolve resistance, potentially reducing the efficacy of the method (Bull & Turelli, 2013; Edenborough et al., 2021; Ritchie et al., 2018). Sequencing of *Ae. aegypti* up to 8 years post release in Cairns, Australia, revealed that host genomes remained mostly stable (Lau et al., 2021), however further work is needed to understand if this translates to other populations. Previous laboratory studies using artificial selection have suggested *Ae. aegypti* genetic variation can impact the strength of *w*Mel-mediated virus blocking (Ford et al., 2019), indicating the need to monitor changes to this genome in addition to critical *Wolbachia*-induced phenotypes over time. The evolution of viruses in *w*Mel-infected hosts has not been extensively studied. RNA viruses are known to have a high mutation rate but are also exposed to strong purifying selection, in particular, as arboviruses must replicate in both human and insect hosts. Multiple lab evolution studies have shown that DENV and Drosophila C virus did not evolve to evade inhibition by *Wolbachia* (Koh et al., 2019; Martinez et al., 2019). However, a recent study showed that selective pressure from *w*Mel resulted in positive selection on DENV (Thi Hue Kien et al., 2023). The viral kinetics of the positively selected SNP have not been evaluated in *w*Mel-infected mosquitoes, so it is unclear whether this change results in evasion of *w*Mel inhibition. Overall, these data highlight the importance of monitoring all three aspects of this tripartite interaction.

Here, we report a multi-country sequencing analysis of 100 individual *w*Mel genomes in *Ae. aegypti*. We assessed what changes occurred to the *w*Mel genome after introgression into three global *Ae. aegypti* populations, and post-field release up to six years. We show *w*Mel genome stability in Colombian, Indonesian, and Vietnamese *Ae. aegypti* field populations, finding limited genomic changes. Despite the different geographical locations, these *w*Mel populations show little divergence since introduction, indicating that *w*Mel has not undergone selection in response to local, country-specific selective pressures. This indicates the use of *w*Mel as a biocontrol agent in novel global *Ae. aegypti* populations has had little effect on the *w*Mel genome in the timeframe of this study. These findings provide reassurance toward the potential of the endosymbiont *w*Mel to remain a successful public-health biocontrol method against arboviruses globally.

## Acknowledgements

We would like to thank I’ah Donovan-Banfield for technical assistance. This research/work was supported by **Monash eResearch** capabilities, including HPC (M3/MASSIVE).

