## Supplemental Data for "High genomic stability of *w*Mel *Wolbachia* after introgression in three geographically distinct *Aedes aegypti* populations"

**Supplementary Material**

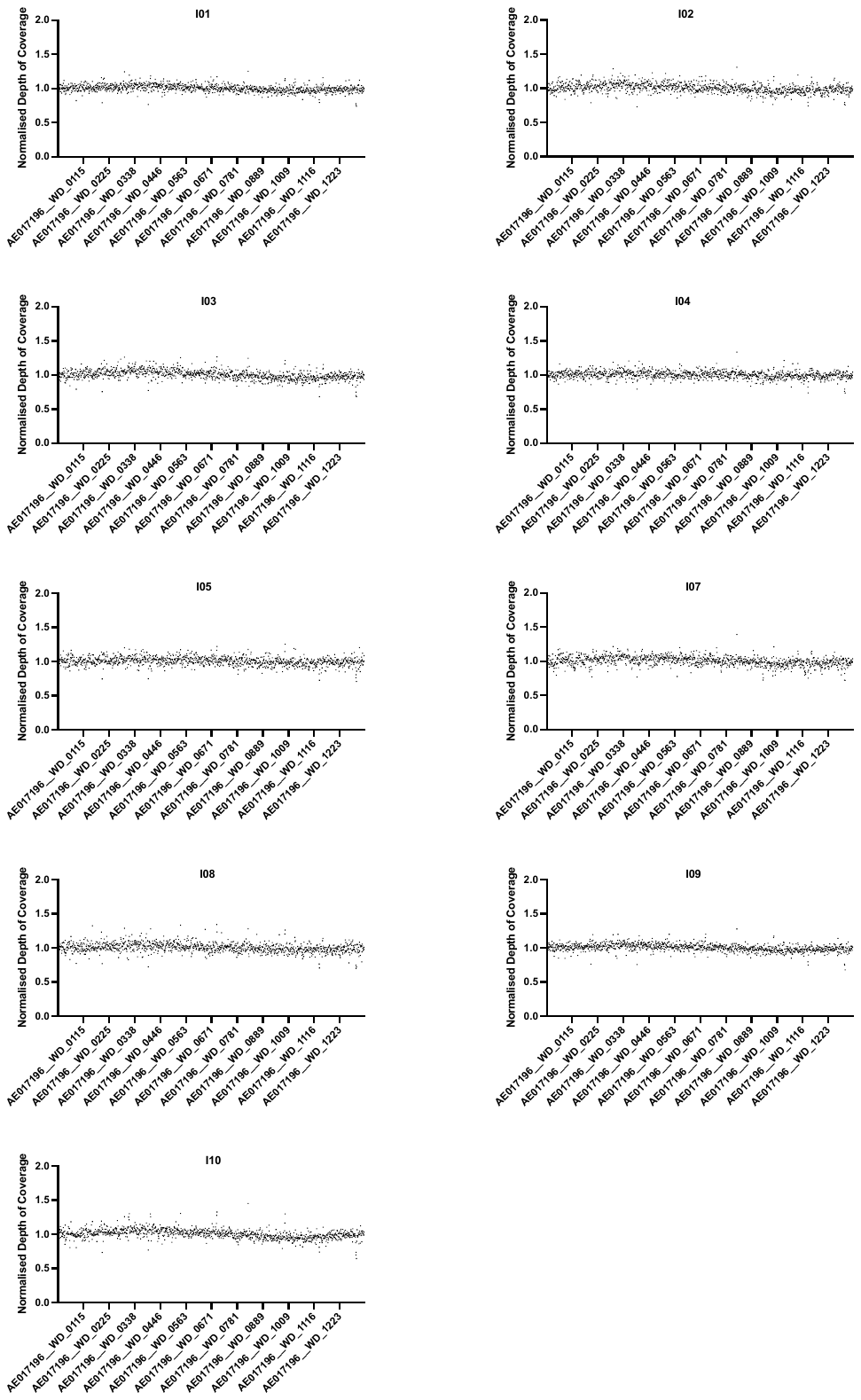
A)

B)

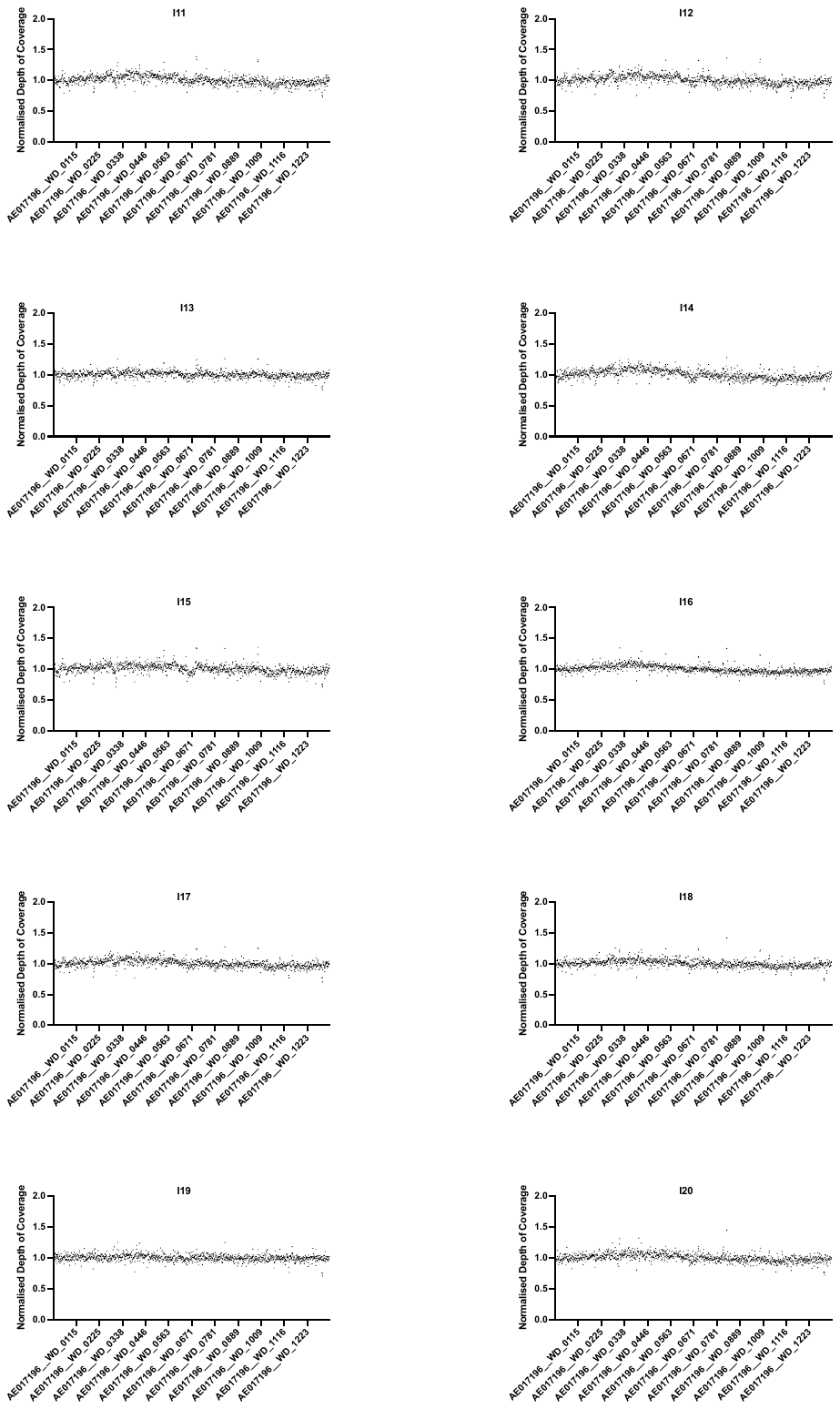

B cont.)

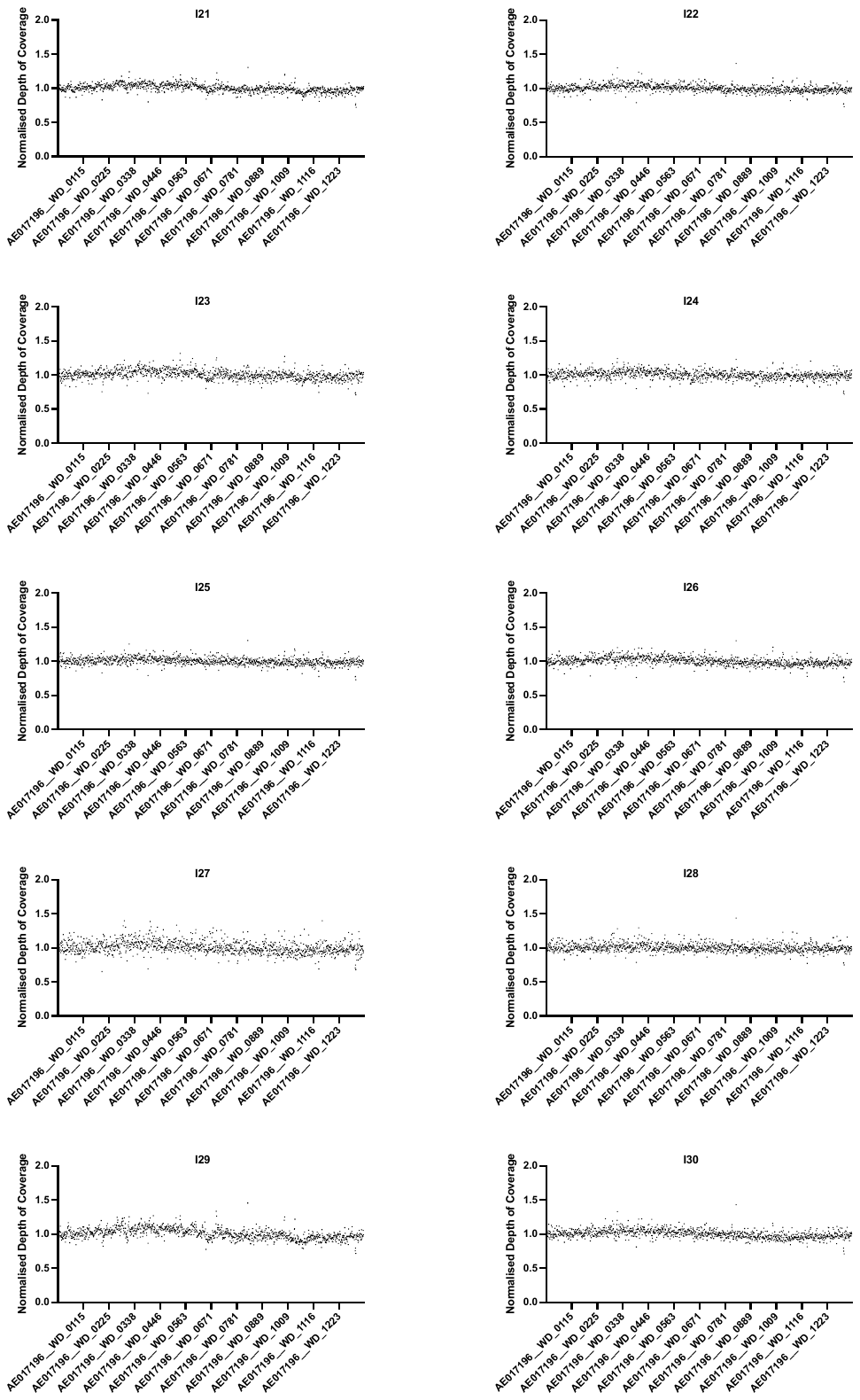

B cont.)

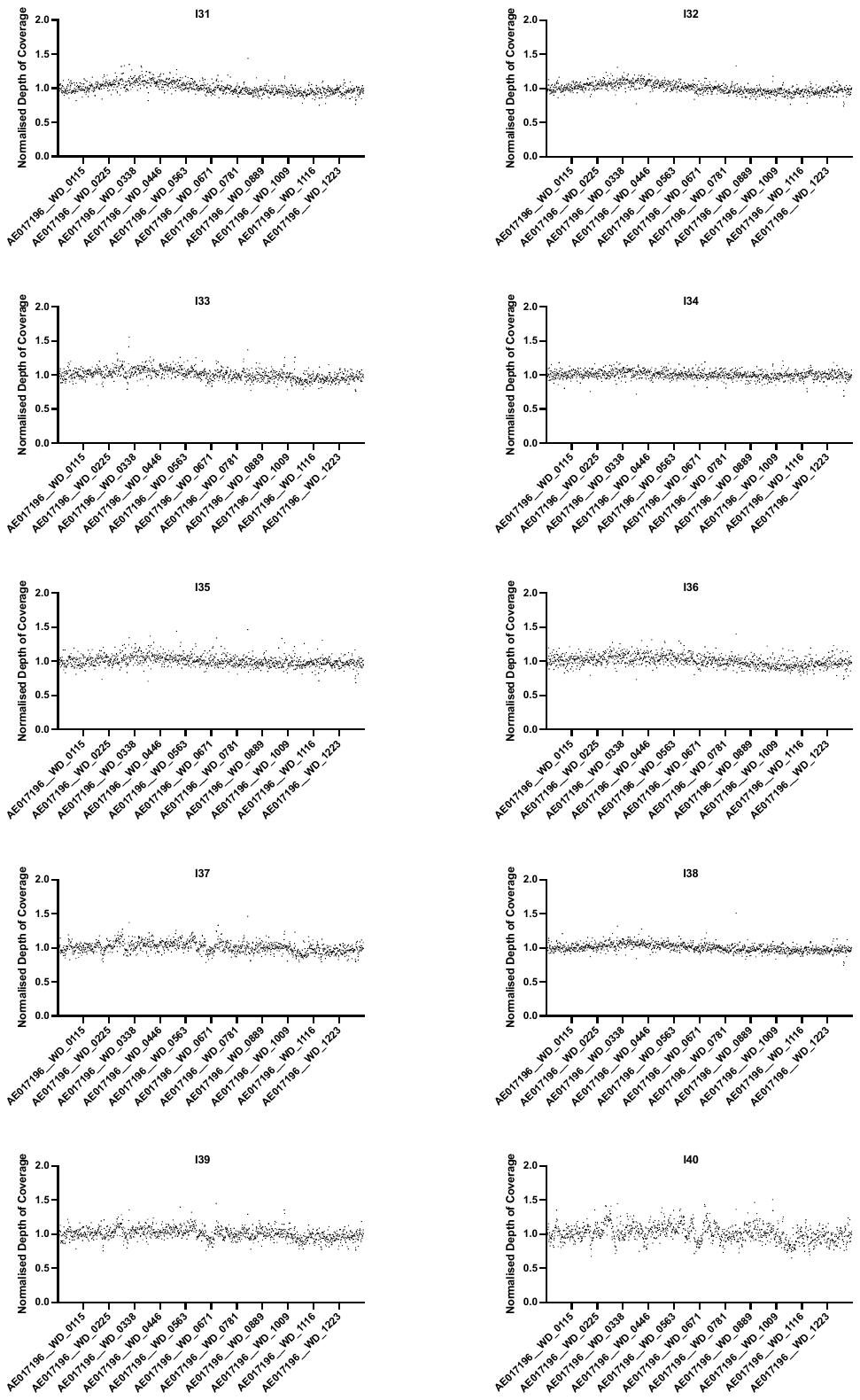

C)

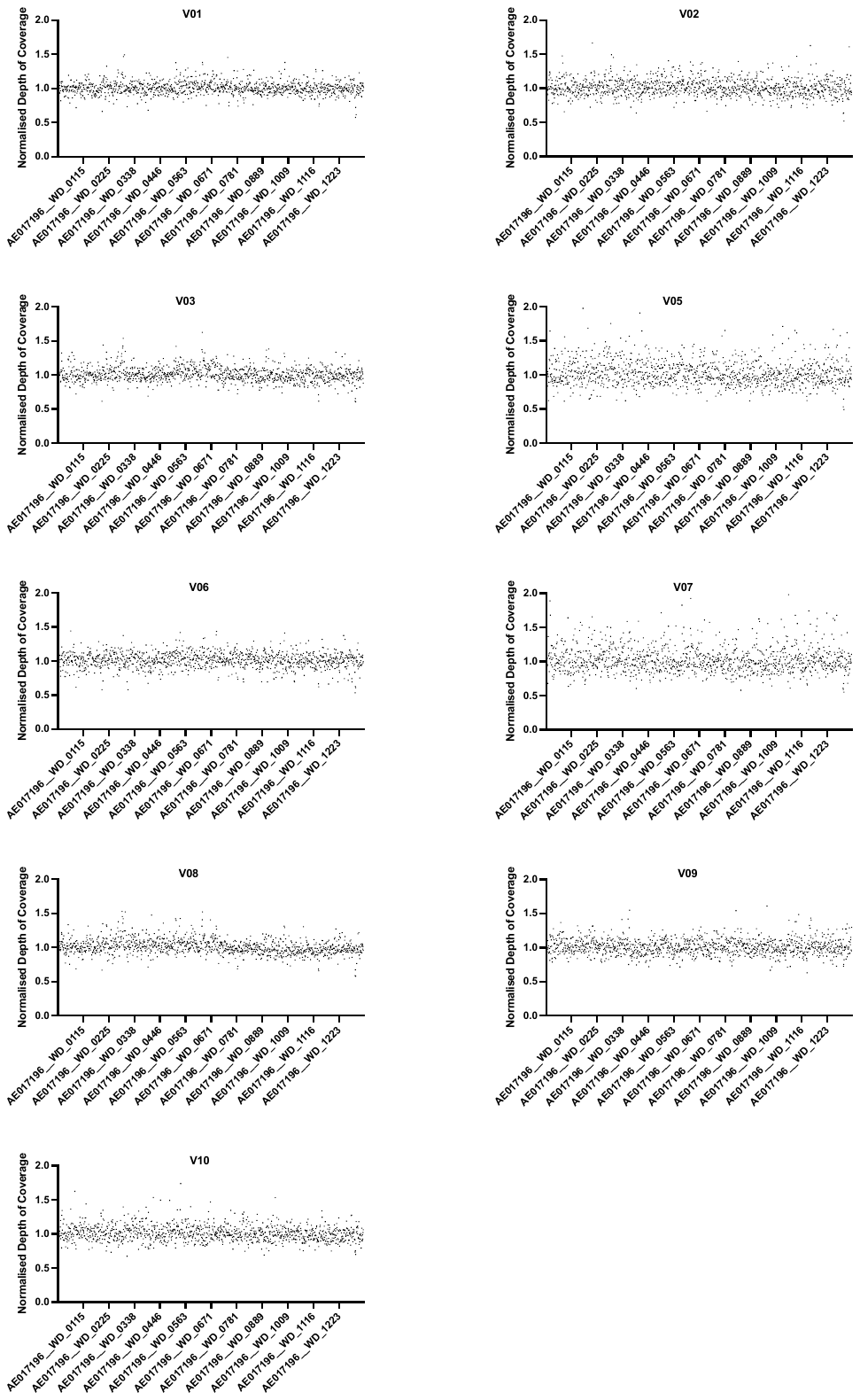

D)

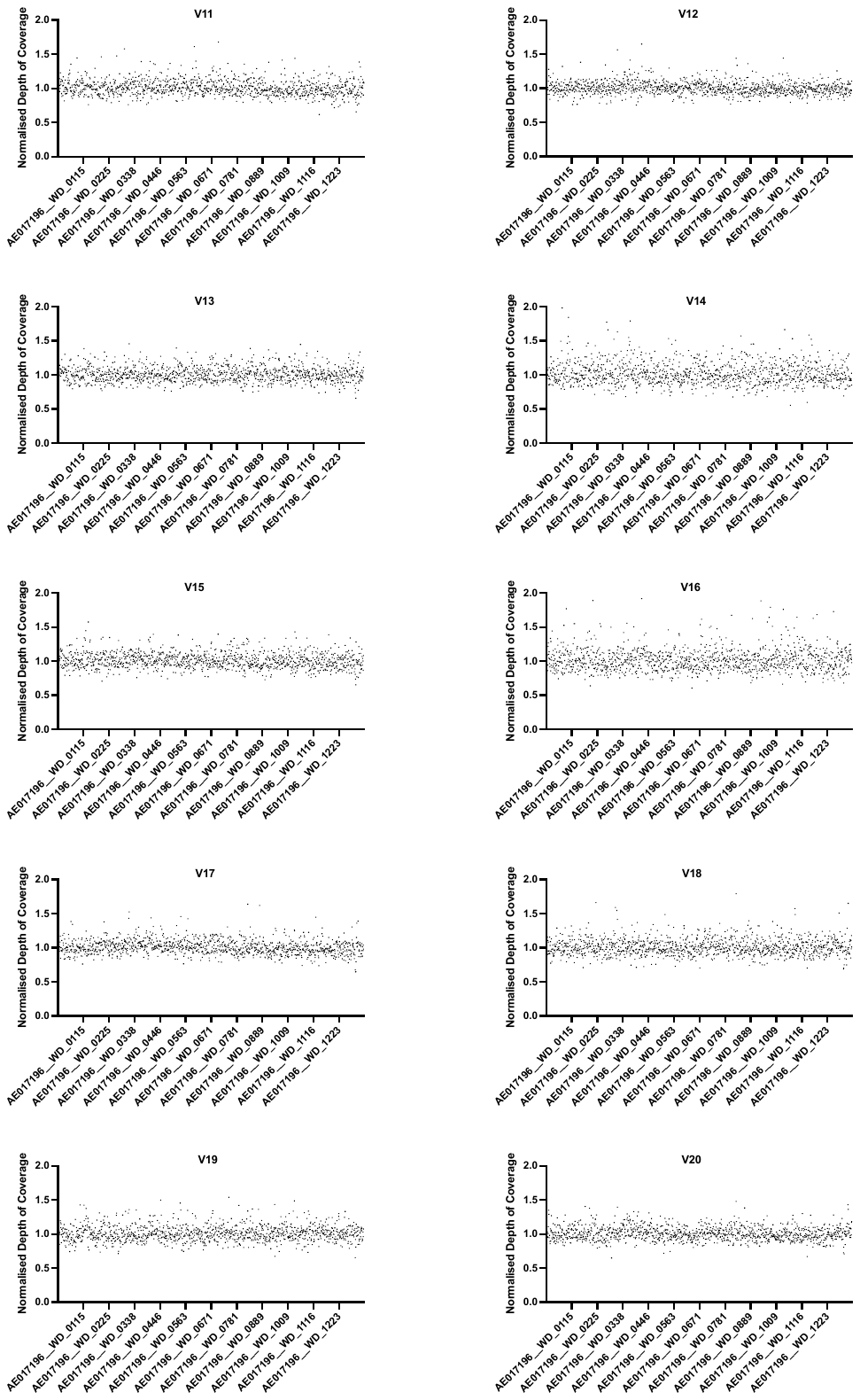

D cont.)

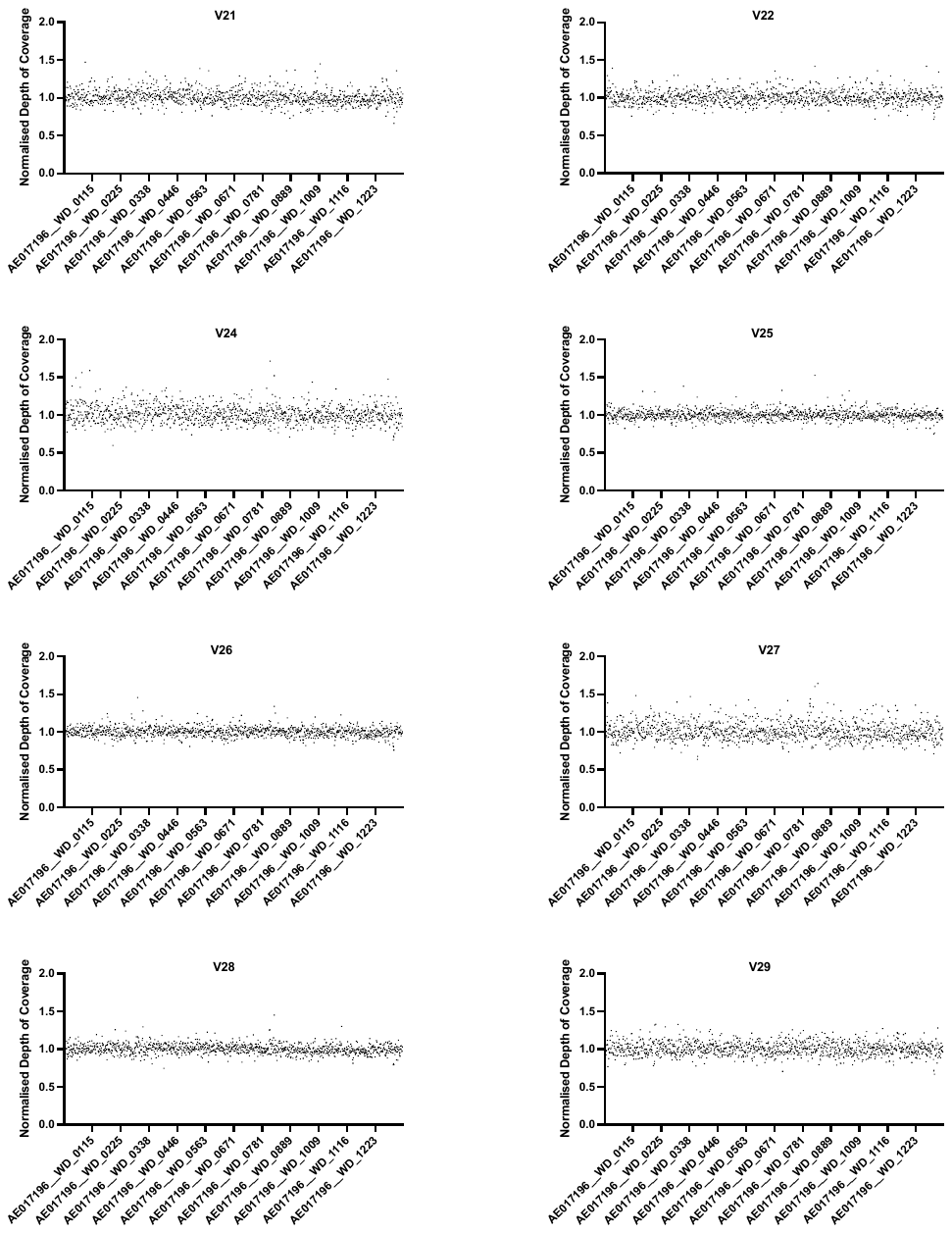

E)

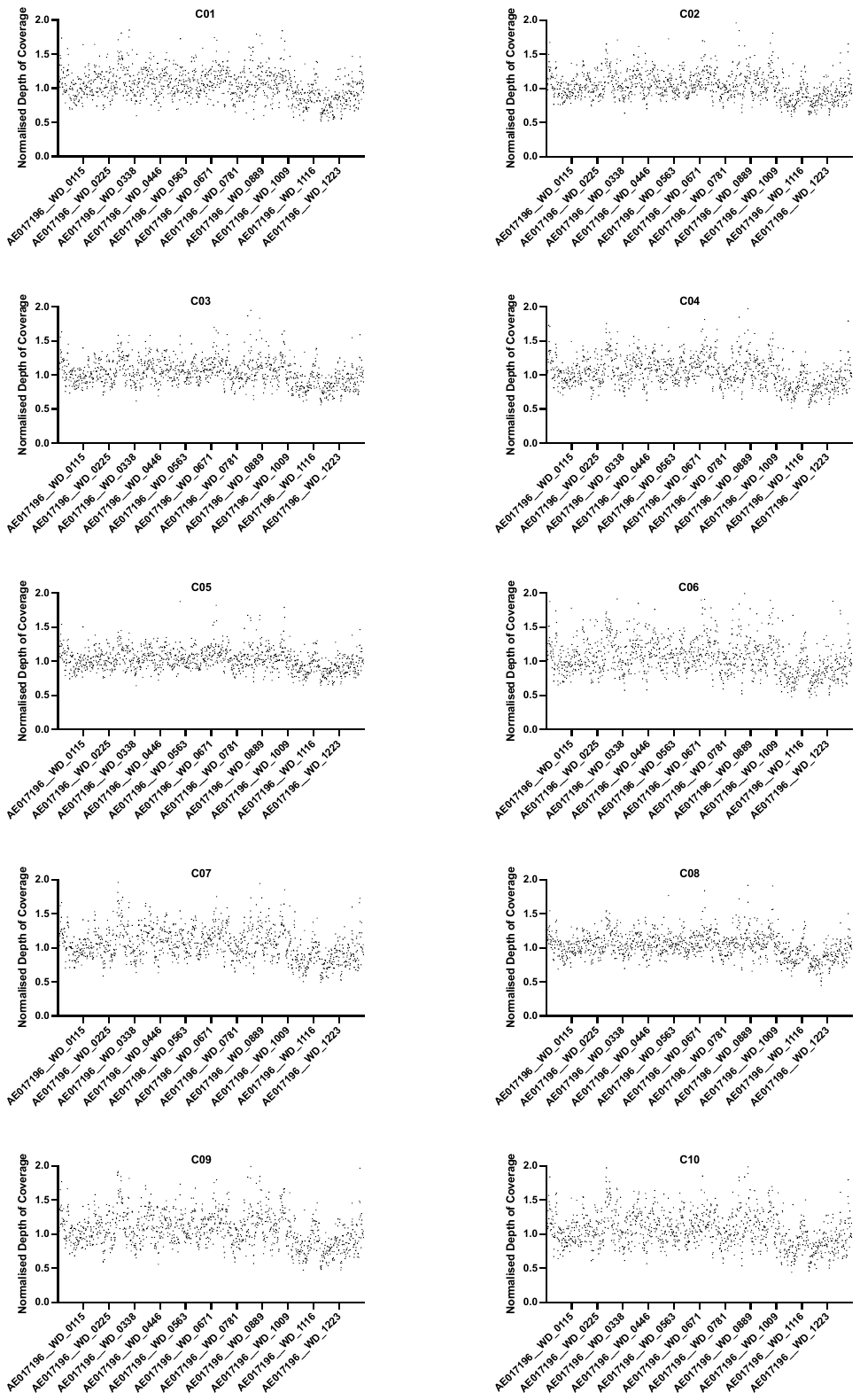

F)

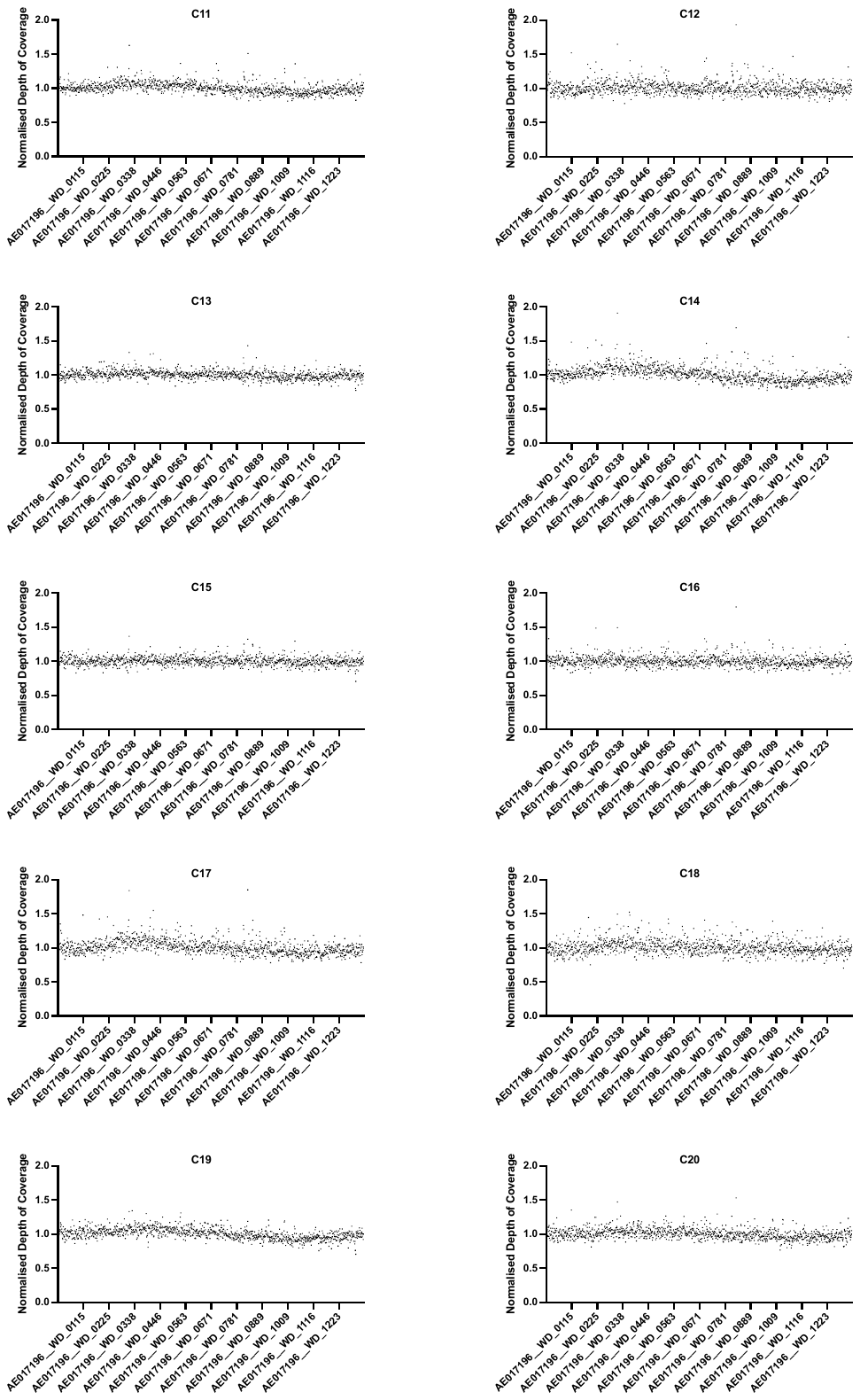

F cont.)

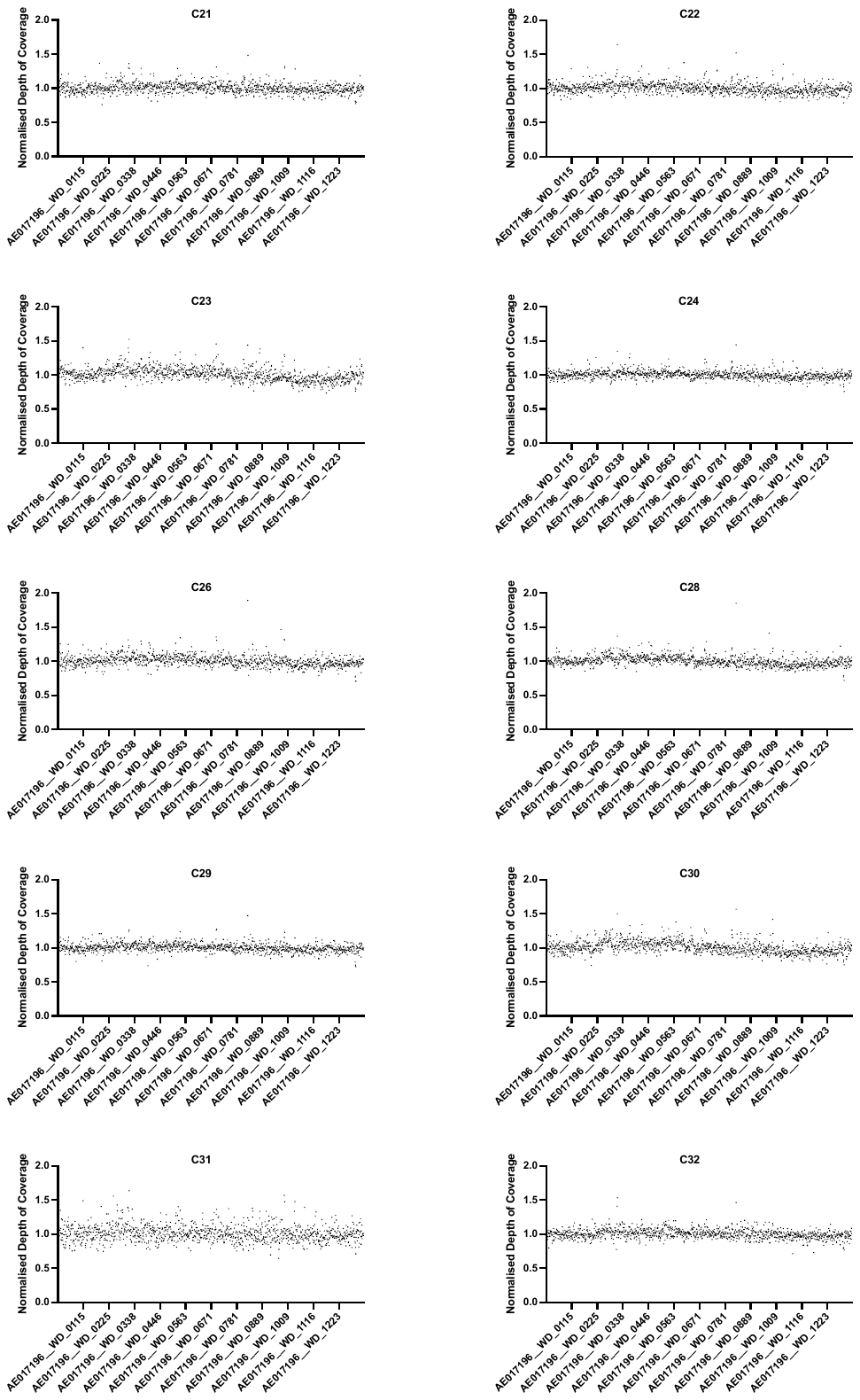

F cont.)

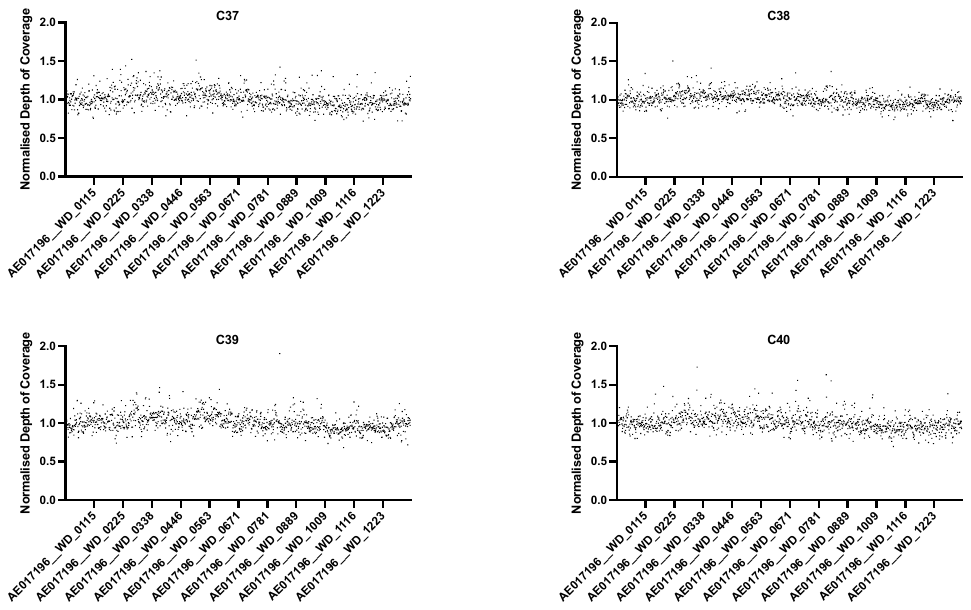

**Supplementary Figure 1. Normalised sequencing depth of coverage.** The distribution of coverage is plotted for all genes across each genome from A) Indonesia 2015, B) Indonesia 2020, C) Vietnam 2015, D) Vietnam 2019, E) Colombia 2015, F) Colombia 2019. Each data point represents the normalised gene depth of coverage for that gene. Normalisation was calculated by dividing the depth of coverage for each gene by the mean depth of coverage for the whole genome.

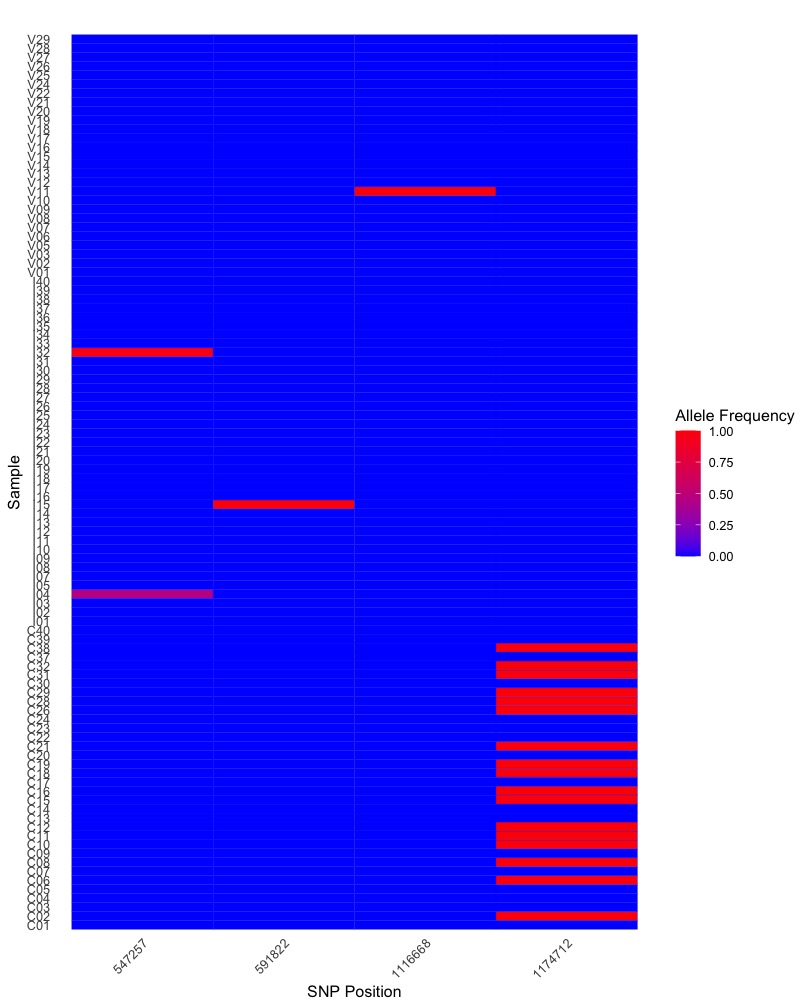

**Supplementary Figure 2: Identified SNP frequencies across all genomes sequenced.** Heatmap showing proportion of reads that mapped to the alternate SNP alleles for each genome sequenced as identified by LoFreq V2.1.3.1. Rows correspond to individual samples, and columns correspond to the four identified SNPs.

**Supplementary Table 1. Low frequency SNPs identified per genome.** Number of low-frequency SNPs identified per genome at frequencies 0-5%, 5-10%, 10-20%, and >20%, and total.

| Population | Genome | Total | 0-5% | 5-10% | 10-20% | >20% |
| --- | --- | --- | --- | --- | --- | --- |
| Vietnam 2015 | V1 | 1519 | 1141 | 315 | 60 | 3 |
| Vietnam 2015 | V2 | 243 | 0 | 11 | 180 | 52 |
| Vietnam 2015 | V3 | 551 | 8 | 249 | 239 | 55 |
| Vietnam 2015 | V5 | 154 | 0 | 0 | 38 | 116 |
| Vietnam 2015 | V6 | 832 | 22 | 324 | 369 | 117 |
| Vietnam 2015 | V7 | 94 | 0 | 0 | 27 | 67 |
| Vietnam 2015 | V8 | 602 | 2 | 278 | 285 | 37 |
| Vietnam 2015 | V9 | 38 | 0 | 7 | 29 | 2 |
| Vietnam 2015 | V10 | 36 | 0 | 2 | 30 | 4 |
| Vietnam 2019 | V11 | 37 | 0 | 6 | 29 | 2 |
| Vietnam 2019 | V12 | 115 | 2 | 55 | 57 | 1 |
| Vietnam 2019 | V13 | 41 | 0 | 1 | 35 | 5 |
| Vietnam 2019 | V14 | 45 | 0 | 0 | 11 | 34 |
| Vietnam 2019 | V15 | 51 | 0 | 3 | 38 | 10 |
| Vietnam 2019 | V16 | 33 | 0 | 0 | 13 | 20 |
| Vietnam 2019 | V17 | 68 | 0 | 37 | 29 | 2 |
| Vietnam 2019 | V18 | 47 | 0 | 0 | 33 | 14 |
| Vietnam 2019 | V19 | 77 | 0 | 5 | 55 | 17 |
| Vietnam 2019 | V20 | 76 | 0 | 18 | 53 | 5 |
| Vietnam 2019 | V21 | 160 | 0 | 48 | 98 | 14 |
| Vietnam 2019 | V22 | 107 | 2 | 49 | 48 | 8 |
| Vietnam 2019 | V24 | 85 | 0 | 3 | 72 | 10 |
| Vietnam 2019 | V25 | 194 | 52 | 119 | 22 | 1 |
| Vietnam 2019 | V26 | 197 | 25 | 139 | 32 | 1 |
| Vietnam 2019 | V27 | 33 | 0 | 4 | 27 | 2 |
| Vietnam 2019 | V28 | 405 | 43 | 161 | 64 | 137 |
| Vietnam 2019 | V29 | 197 | 3 | 105 | 85 | 4 |
| Colombia 2015 | C1 | 515 | 19 | 299 | 182 | 15 |
| Colombia 2015 | C2 | 858 | 356 | 401 | 97 | 4 |
| Colombia 2015 | C3 | 379 | 129 | 212 | 37 | 1 |
| Colombia 2015 | C4 | 430 | 144 | 249 | 36 | 1 |
| Colombia 2015 | C5 | 1556 | 826 | 561 | 160 | 9 |
| Colombia 2015 | C6 | 791 | 11 | 310 | 313 | 157 |
| Colombia 2015 | C7 | 743 | 138 | 424 | 170 | 11 |
| Colombia 2015 | C8 | 1109 | 428 | 506 | 166 | 9 |
| Colombia 2015 | C9 | 644 | 198 | 310 | 130 | 6 |
| Colombia 2015 | C10 | 1255 | 363 | 560 | 284 | 48 |
| Colombia 2019 | C11 | 202 | 85 | 94 | 18 | 5 |
| Colombia 2019 | C12 | 57 | 2 | 45 | 8 | 2 |
| Colombia 2019 | C13 | 360 | 197 | 153 | 9 | 1 |
| Colombia 2019 | C14 | 372 | 251 | 93 | 27 | 1 |
| Colombia 2019 | C15 | 247 | 114 | 115 | 16 | 2 |
| Colombia 2019 | C16 | 108 | 15 | 75 | 16 | 2 |
| Colombia 2019 | C17 | 196 | 54 | 118 | 23 | 1 |
| Colombia 2019 | C18 | 115 | 0 | 50 | 61 | 4 |
| Colombia 2019 | C19 | 129 | 5 | 72 | 48 | 4 |
| Colombia 2019 | C20 | 238 | 32 | 167 | 38 | 1 |
| Colombia 2019 | C21 | 356 | 93 | 226 | 35 | 2 |
| Colombia 2019 | C22 | 123 | 9 | 101 | 12 | 1 |
| Colombia 2019 | C23 | 216 | 17 | 135 | 60 | 4 |
| Colombia 2019 | C24 | 358 | 158 | 168 | 31 | 1 |
| Colombia 2019 | C26 | 677 | 172 | 410 | 91 | 4 |
| Colombia 2019 | C28 | 498 | 253 | 208 | 34 | 3 |
| Colombia 2019 | C29 | 759 | 402 | 297 | 54 | 6 |
| Colombia 2019 | C30 | 207 | 49 | 123 | 34 | 1 |
| Colombia 2019 | C31 | 358 | 5 | 186 | 152 | 15 |
| Colombia 2019 | C32 | 117 | 33 | 77 | 5 | 2 |
| Colombia 2019 | C37 | 70 | 0 | 27 | 40 | 3 |
| Colombia 2019 | C38 | 52 | 0 | 29 | 21 | 2 |
| Colombia 2019 | C39 | 104 | 0 | 64 | 37 | 3 |
| Colombia 2019 | C40 | 85 | 0 | 33 | 46 | 6 |
| Indonesia 2015 | I1 | 1519 | 1141 | 315 | 60 | 3 |
| Indonesia 2015 | I2 | 1685 | 1241 | 364 | 76 | 4 |
| Indonesia 2015 | I3 | 738 | 227 | 401 | 103 | 7 |
| Indonesia 2015 | I4 | 1044 | 753 | 246 | 41 | 4 |
| Indonesia 2015 | I5 | 463 | 95 | 256 | 106 | 6 |
| Indonesia 2015 | I7 | 607 | 317 | 227 | 61 | 2 |
| Indonesia 2015 | I8 | 784 | 265 | 381 | 126 | 12 |
| Indonesia 2015 | I9 | 1637 | 1190 | 375 | 69 | 3 |
| Indonesia 2015 | I10 | 803 | 295 | 371 | 130 | 7 |
| Indonesia 2020 | I11 | 1201 | 719 | 374 | 102 | 6 |
| Indonesia 2020 | I12 | 1158 | 536 | 497 | 116 | 9 |
| Indonesia 2020 | I13 | 1112 | 741 | 316 | 54 | 1 |
| Indonesia 2020 | I14 | 684 | 433 | 216 | 34 | 1 |
| Indonesia 2020 | I15 | 1097 | 581 | 410 | 99 | 7 |
| Indonesia 2020 | I16 | 1233 | 917 | 266 | 48 | 2 |
| Indonesia 2020 | I17 | 1787 | 1228 | 462 | 93 | 4 |
| Indonesia 2020 | I18 | 1764 | 1345 | 346 | 70 | 3 |
| Indonesia 2020 | I19 | 1304 | 1065 | 200 | 38 | 1 |
| Indonesia 2020 | I20 | 786 | 499 | 235 | 49 | 3 |
| Indonesia 2020 | I21 | 1463 | 1122 | 281 | 58 | 2 |
| Indonesia 2020 | I22 | 1495 | 1237 | 223 | 35 | 0 |
| Indonesia 2020 | I23 | 883 | 384 | 405 | 89 | 5 |
| Indonesia 2020 | I24 | 909 | 599 | 232 | 75 | 3 |
| Indonesia 2020 | I25 | 1262 | 1005 | 224 | 31 | 2 |
| Indonesia 2020 | I26 | 1875 | 1528 | 295 | 50 | 2 |
| Indonesia 2020 | I27 | 1700 | 971 | 572 | 145 | 12 |
| Indonesia 2020 | I28 | 1688 | 1513 | 146 | 27 | 2 |
| Indonesia 2020 | I29 | 1008 | 592 | 291 | 82 | 43 |
| Indonesia 2020 | I30 | 560 | 345 | 198 | 16 | 1 |
| Indonesia 2020 | I31 | 461 | 145 | 247 | 67 | 2 |
| Indonesia 2020 | I32 | 576 | 283 | 235 | 55 | 3 |
| Indonesia 2020 | I33 | 332 | 78 | 195 | 56 | 3 |
| Indonesia 2020 | I34 | 1685 | 1241 | 364 | 76 | 4 |
| Indonesia 2020 | I35 | 1021 | 400 | 466 | 147 | 8 |
| Indonesia 2020 | I36 | 359 | 50 | 233 | 73 | 3 |
| Indonesia 2020 | I37 | 505 | 225 | 235 | 44 | 1 |
| Indonesia 2020 | I38 | 395 | 252 | 124 | 18 | 1 |
| Indonesia 2020 | I39 | 484 | 100 | 224 | 145 | 15 |
| Indonesia 2020 | I40 | 325 | 44 | 209 | 70 | 2 |

**Supplementary Table 2 Mean number of low-frequency variants, mean *w*Mel density in samples that were sequenced, and mean sequence depth per genome of each population sequenced.**

| **Population** | **Mean number of variants at frequencies 10-20% per genome** | **Mean number of variants at frequencies >20% per genome** | **Mean *w*Mel density** | **Mean Sequence depth** |
| --- | --- | --- | --- | --- |
| **Indonesia 2015** | 85.78 | 5.33 | 23.5 | 807 |
| **Indonesia 2020** | 68.73 | 5.03 | 25.3 | 858.3 |
| **Vietnam 2015** | 139.67 | 50.33 | 4.01 | 54.8 |
| **Vietnam 2019** | 44.50 | 15.94 | 4.14 | 65.0 |
| **Colombia 2015** | 157.5 | 26.10 | 11.84 | 338.4 |
| **Colombia 2019** | 38.17 | 3.17 | 13.59 | 230.8 |

**Supplementary Table 3. List of IS queries identified to have greater than 80% similarity to the *w*Mel reference genome.**

| IS Element | Family | Group |
| --- | --- | --- |
| ISCaa8 | IS5 | IS903 |
| ISWen1 | IS4 | IS231 |
| isWen2 | IS110 |  |
| iswen3 | IS66 | ISBst12 |
| iswosp2 | IS4 | IS50 |
| iswpi1 | IS5 | IS1031 |
| iswpi11 | IS630 |  |
| iswpi12 | IS110 |  |
| iswpi14 | IS110 |  |
| iswpi15 | IS256 |  |
| iswpi18 | IS4 | IS4 |
| iswpi4 | IS481 |  |
